# Individual performance niches may buffer population responses to climate change in estuarine fishes

**DOI:** 10.1101/2024.02.01.578478

**Authors:** Clara Bellotto, Ashley M. Fowler, David J. Booth

## Abstract

Climate change may impact individual organisms in different ways, a consideration often overshadowed by predominant focus on population effects in studies. We examined three estuarine fish species to determine if individual fish performance, persisted across winter water temperatures. Fish performance at 16°C (current Sydney winter estuarine water temperature) and 20°C (predicted under climate change) with low and high food regimes was assessed using key physiological (growth, aerobic scope, burst speed) and behavioural parameters (foraging activity, boldness, shelter usage, predator escape response). We expected a strong positive relationship between performance at 16°C and 20°C for each parameter, and interactions with food level, however in general this was not found for any species. Relative performance was only maintained across temperatures for a few parameters, such as bite rate, boldness, and shelter response in one species (trumpeter *Pelates sexlineatu*s), with aerobic scope in silver biddy *Gerres subfasciatus*, and boldness in fortescue *Centropogon australis*.

Our results suggest that individuals’ fitness (directly via changes in growth, indirectly via behaviours) will be impacted by climate warming due to differences in relative performance among individuals across water temperatures. Changes in relative performance among individuals may initially compensate for a population-level response, thereby buffering the effects of climate change.

## 1. Background

An organism’s ability to cope with environmental change is affected by its capacity to initiate an appropriate stress response via physiological systems, reallocating energy towards defensive mechanisms, and making behavioural adjustments to deal with or evade the environmental threat (1). For instance, changes in water temperature directly modify fish physiological conditions, growth and developmental rates, metabolism, muscle and cardiovascular function, swimming ability, behaviour and reproductive performance (2–4). Energy metabolism, as measured by oxygen consumption and aerobic scope (4, 5), plays a pivotal role in a fish’s physiological fitness. Optimal fitness is achieved within a specific temperature range, with performance following a bell-shaped curve where performance is optimized at intermediate temperatures (6). Both the shape of the performance curve and the optimal temperature differ among species (7, 8). Temperature also affects biochemical efficiencies, leading to potential growth and population responses, alongside heightened oxidative stress (9). Moreover, rising temperatures can alter the balance between metabolic rates, impacting aerobic scope, cardiac scope, and overall fish growth and survival (4). While increased temperature often corresponds to improved growth rates and survival, these benefits may diminish when temperatures exceed the optimum (10, 11).

There has been a growing emphasis on the contribution of physiology to evolutionary processes and the influence of physiological traits on life-history trade-offs (Metcalfe et al. 2016). This highlights the need to examine the individual variability in physiological characteristics because individual differences drive heritable variation, thus population change (12). This approach to ecophysiology focuses on evaluating the causes and ecological implications of individual differences in physiological traits instead of considering them as variance around a population mean, (13). For instance, considering population performance across temperatures for a specific metric (Figure 1A), the ‘error bar’ representing variation within the population could, for example, stem from different scenarios. In one case (Figure 1B), individual performance trends align consistently with the population trend. In another scenario (Figure 1C), individual trends across temperatures differ, yet collectively result in the same mean and error value for the populations. The initial reordering of individuals in scenario C may temporarily delay a population effect as temperature increases, because individuals may exchange performance ranks without affecting the population mean or variance.

**Figure 1.**
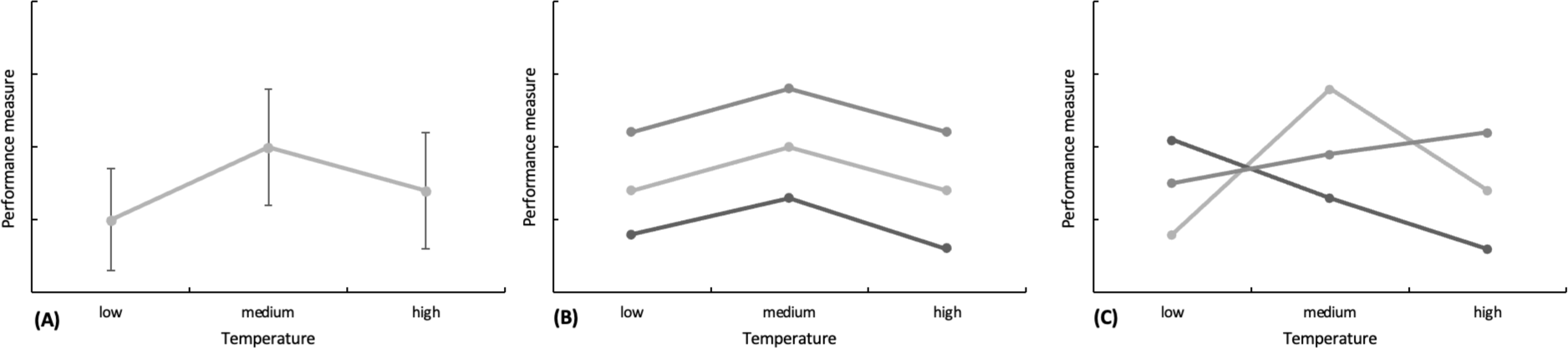
(**A**) Hypothetical mean population performance across different temperatures (± SEM). (**B**) Individual performance from the individuals that comprise population from Figure 1A, where each line represents one individual. Average population performance from figure A results from the three individual performances averaged with consistent trend across individuals. (**C**) Individual performance from the individuals that comprise population from figure A, where each line represents one individual. Average population performance from figure A results from the three individual performances averaged with different trend across individuals.

**Figure 2.**
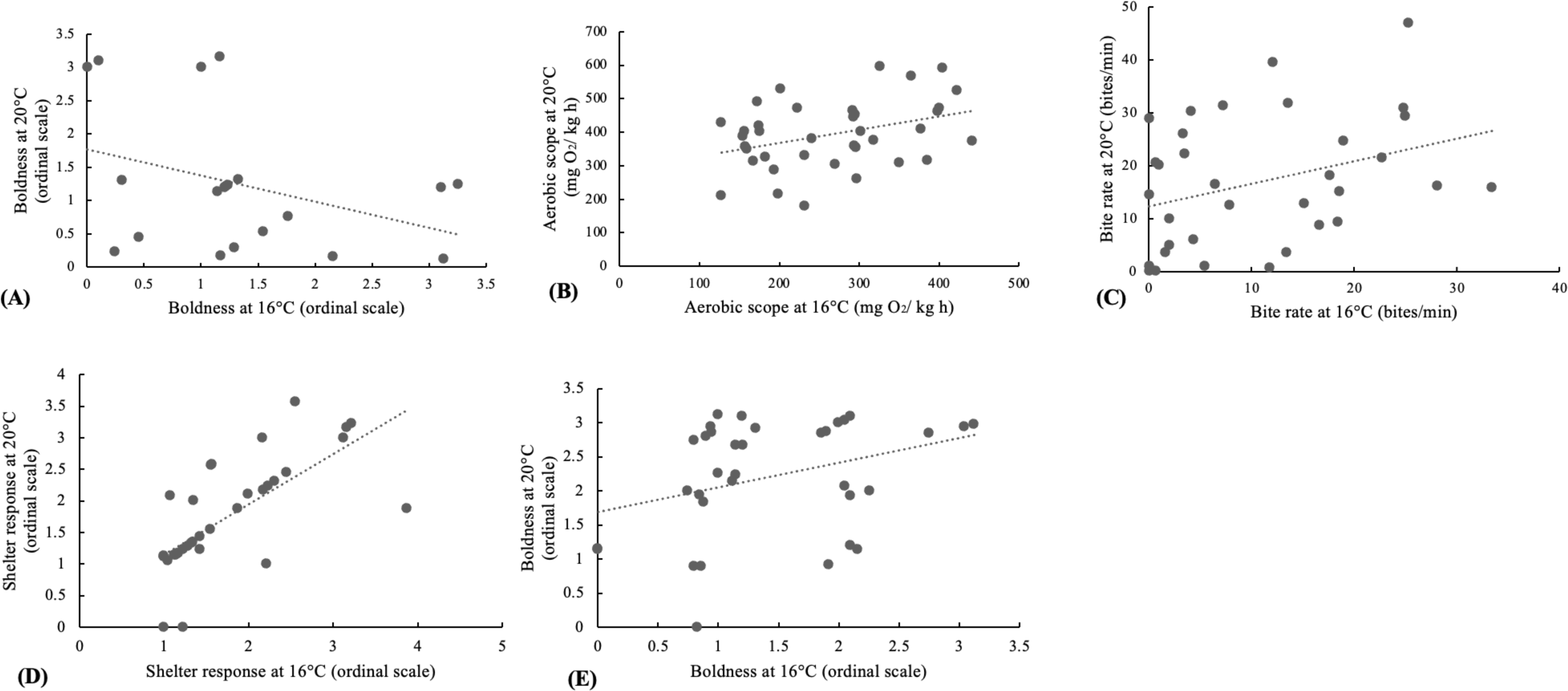
Regressions (p-value <0.05) for some performance metrics between individual performance at 16°C vs individual performance at 20°C across species. **(A)** *Centropogon australis* boldness (ordinal scale; points were jittered for graphing purposes); **(B)** *Gerres subfasciatus* aerobic scope (mg O_2_/ kg h); **(C)** *P. sexlineatus* bite rate (bites/min); **(D)** *P. sexlineatus* shelter response (ordinal scale; points were jittered for graphing purposes); **(E)** *P. sexlineatus* boldness (ordinal scale; points were jittered for graphing purposes).

Only a limited number of studies have assessed fish individual performance across different environmental conditions. These studies suggest that physiological performance and personality traits such as activity, aggressiveness, and boldness can be preserved in individuals that are exposed to changing environmental conditions (14–19). This behavioural consistency is vital as it can affect social hierarchies and, thus, population dynamics as differences in traits like activity level, escape response, and boldness influence foraging efficiency and predator interactions (20). For instance, boldness is an essential factor that determines an individual’s position within a social network as either dominant or subordinate in both three-spined sticklebacks (*Gasterosteus aculeatus*) and guppies (*Poecilia reticulate*); this will ultimately affect reproductive success and genetic inheritance which could thus vary with changes in boldness dictated by environmental conditions (21, 22). While bold behaviours can enhance certain aspects of an individual’s life, such as dominance, migration, foraging, and reproduction, they may also entail increased predation risks, potentially affecting long-term survival (23). Rising temperatures often correlate with heightened activity and boldness among fish (24, 25). Moreover, alterations in physiological traits can drive shifts in behaviour (25). High metabolic rates tend to align with increased boldness and activity (26, 27). As temperatures rise, the heightened metabolic demand may lead to more risk-taking behaviours to boost energy intake (28). However, to the authors’ knowledge, no previous study has examined individual performance of fish across a temperature gradient.

Escape response, a critical aspect of fish performance for predator avoidance, can also be influenced by temperature shifts (29). While some species exhibit higher burst speeds and escape responses in warmer conditions, others may display reduced responsiveness (30–32). Shelter usage is another significant performance attribute influencing survival, reproductive success, and energy allocation (33). Balancing the need to avoid predators with the exploration of resources presents trade-offs that shape fish behaviour and, ultimately, influence their reproduction and survival (34). Recognizing these complex interrelationships is pivotal for comprehending the adaptive value of different phenotypes, which can have enduring effects on future generations (13).

The performance functions of individuals can exhibit heritable variation, which means that differences in these traits can be inherited from parents by their offspring across generations (Donelson et al. 2011). As there is a positive relationship between performance and fitness, natural selection will favour the individuals that perform better at the new body temperature experienced compared to the old one (35). However, it is expected that some evolutionary trade-offs will arise that could be manifest in the thermal sensitivity of performance (e.g., a negative relationship between different performance metrics at low vs high temperatures) (36).

Considering climate warming, some individuals performing best at high temperatures might take advantage of the newly created warmer niche. At the same time, intraspecific competition might increase for the other subset of individuals that share the same functional group as their suitable (cooler) thermal niche space shrinks (37).

In light of climate change-induced sea temperature rise, the species that present wide thermo-tolerance and physiological adaptability might be at an advantage over others (38). Estuaries are highly dynamic environments, an interface between freshwater and marine habitats where fluctuations in abiotic factors such as temperature are frequent (39). As such, we would expect estuarine species to be adapted to such a changing environment (40).

Consequently, estuarine species are expected to have high plasticity and thus be a valuable model for impacts of future climate change. The performance of estuarine fish at various temperatures, including winter conditions, remains an understudied area, with potential implications for stock densities and growth potential, particularly in species recruiting during winter (41, 42). In addition, given food limitation that often occurs over winter, how performance relative to temperature interacts with food level is important to evaluate as lower food regimes/deprivation are linked to changes performance (growth, aerobic scope, swimming speed, boldness) in fish (43–45). Furthermore, the epigenome’s susceptibility to environmental influences during early life stages underscores the need for species- and life-stage-specific investigations into temperature’s impacts on fish performance over winter (4, 46).

The south-eastern region of Australia has witnessed significant rises in average ocean temperatures, primarily attributed to the intensification of the East Australian Current (47, 48), with the projection that Australian temperate marine waters will increase by 1.5°C to 3°C by 2070 (49). Recent findings indicate that Australian estuaries - key fish habitats and nurseries are experiencing even faster warming at a rate of 0.2 °C per year (40). Additionally, climate change causes alterations in food resources, potentially leading to reduced availability for certain fish functional groups (37).

This study aimed to test whether individuals of estuarine fishes have individual performance niches within the overall population. Specifically, we aimed to determine the individual performance of juvenile estuarine fish across various physiological and behavioural measures at two different food regimes (high and low food) and temperatures that reflect current and forecasted winter conditions in south-eastern Australia; specifically, 16°C (typical cold winter) and 20°C (projected warming)(50). Three common estuarine species in the region were investigated. We predicted that relative individual performance would be maintained across treatments for all metrics.

## 2. Materials and Methods

### a) **Fish species and collection**

Juvenile fish were obtained using small hand-drawn seine nets from seagrass (*Zostera* sp. and *Posidonia* sp.) in Careel Bay, within in the Pittwater estuary located in the south-eastern part of Australia (33° 3’ 02.8’’ S; 151° 1’ 25.2’’ E). After capture, the fish were transferred to the aquarium facilities at the University of Technology Sydney (UTS).

This study was conducted on juveniles of three temperate estuarine species: the eastern fortescue, *Centropogon australis*, *Scorpaenidae* the common silverbiddy, *Gerres* subfasciatus, *Gerreidae* and the eastern striped trumpeter, *Pelates sexlineatus*, *Tetrapontidae* These specific species were chosen because they were common at the collection site and so a sufficient number of individuals were available.

### **b)** Laboratory husbandry and acclimation

The captured fish were initially housed in groups of 10 to 15 in 40-liter tanks, where they were kept at the ambient temperature of the Sydney estuary at the time of capture for two days (16°C - 20°C). Subsequently, the fish were separated and individually housed in 15-liter tanks filled with natural seawater at a salinity of 35 ppt, sourced from the holding tanks within the UTS facilities (salinities at the field site ranged from 34-35ppt). To acclimatize the fish to the desired temperatures of either 16°C or 20°C, a gradual daily temperature change of 0.5°C was implemented using 25 W heaters. This approach ensured a suitable rate of acclimatization while minimizing stress on the fish, as outlined by (51).

The duration of the acclimatization process ranged from 1 to 7 days, depending on the temperature observed at the collection site and the targeted temperature for the treatment as fish species were collected in different winter months (16°C for *C. australis*, 18°C for *G. subfasciatus* and 20°C for *P.sexlineatus*). The laboratory room air temperature was maintained at 14°C, with lighting on a 12-hour light and 12-hour dark cycle. The fish were fed twice daily with New Life Spectrum® fish pellets. In the tanks, two pieces of polyvinyl chloride (PVC) pipes were introduced as shelter, along with sections of black foam board placed on the tank’s exterior to partially obscure the view and minimize external disturbances, following the method detailed by O’Connor and Booth (52). To maintain water quality, debris such as uneaten food and feces were removed through siphoning, during a daily 30% water change. Additionally, water parameters such as pH, dissolved oxygen, and salinity were monitored every two days using a multi-sensor probe, while water temperature was assessed daily. The health and behaviour of the fish were visually inspected and documented on a daily basis, following the protocol established by Djurichkovic, Donelson (53). The fish remained in the laboratory setting for approximately nine weeks, and upon conclusion of the experimental period, all fish were humanely euthanized using an ice slurry method (54).

### **c)** Experimental protocols

Experimental protocols are shown in Supplementary Figure 1. The performance of fish was evaluated across a number of metrics: metabolic rate, growth, burst speed, foraging behaviour, boldness, shelter use, and escape response (e.g. see Clark *et al*., 2013; O’Connor and Booth, 2021) . Bite rate, burst speed, shelter response, boldness, and escape response were also monitored as important behavioural aspects that shed light on fish performance within their environmental and social contexts (56).

Fish length and mass were measured, and the fish were randomly assigned to one of four treatment groups and placed in individual tanks. All groups (with fishes housed individually) underwent exposure to two different temperatures (16°C and 20°C). However, the sequence in which these temperatures were presented was alternated to account for any potential order-related effects. Within the distinct food regime groups, one group was exposed to temperatures in a descending order (from 20°C to 16°C), while the other group experienced temperatures in an ascending order (from 16°C to 20°C) (Supplementary Figure 1B) to account for order of treatment presentation. Two of the groups were subjected to low food regimes, while the other two experienced high food regimes (Supplementary Figure 1B). If different size classes were present, efforts were made to evenly distribute them across the groups. Fish in the high food treatment were provided with 1% of their body weight in fish pellets twice daily, while those in the low food treatment received 0.5% of their body weight in fish pellets twice a day, following the protocol established by Donelson, Munday (57) and considering that the normal feeding ratio for fish is approximately 0.7% of their body weight in dry food (58). These feeding regimes were consistently maintained throughout the experiments. In the case of the *C. australis* experiment, food regimes were not assessed, and the fish were divided into only two groups based on temperature, with fish being fed *ad libitum,* noting feeding was minimal (Supplementary Figure 1A).

Once the fish reached the initially-required temperature (either 16°C or 20°C), their total length and wet mass were measured. Subsequently, they were exposed to the corresponding temperature treatment for ten days, during which their performance metrics were assessed. Following this ten-day period, the fish were acclimated to the final temperature (20°C or 16°C), and their total length and wet mass were measured once again. They were then exposed to the corresponding temperature treatment for another ten days, after which their performance metrics were once more evaluated, and then the fish were euthanised for otolith analysis.

Each temperature group required approximately three days to complete behavioural assessments and an additional three days for metabolic rate measurements. The selection of individual fish for testing was carried out randomly. For the behavioural tests, fish were transferred to a test tank at the same water temperature, featuring a gridded background with 0.5 cm grids. Observations were recorded using a GoPro® Hero 6 positioned in front of and at the same level as the test tank’s base, capturing videos at a frame rate of 120 frames per second. The fish were allowed a 10-minute acclimation period in the test tank before the commencement of various behavioural tests, each lasting for a 3-minute observation period, with an additional 2-minute acclimation period between tests, following the methodology of Djurichkovic, Donelson (53).

### d) Performance metrics

i. **Fish growth** was quantified by tracking changes in the fish’s total length and wet mass, a method in accordance with the approach described by O’Connor and Booth (52). To estimate the somatic growth rate, we calculated instantaneous growth rate (*G_INST_*).

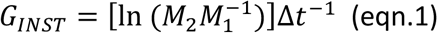

with the mass of each individual fish at the experiment’s outset (*M_1_*) and its mass at the conclusion of the experimental period (*M_2_*), which spanned a specified number of days (t) following equation 1 (59):

We measured changes in the fish’s total length over time, denoted as *TL_c_* by comparing the total length at the experiment’s commencement (*TL_c_*) with that at its conclusion (*TL_2_*), which spanned a specified number of days (t). This assessment was conducted using equation 2:

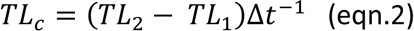

(ii) **Foraging performance** of the fish by measuring their **bite rate**, which represents the total number of feedings strikes they executed per minute. To initiate feeding, a siphon introduced approximately 1.5 grams of fish pellets into the test tank. Following the removal of the siphon, we recorded the bite rate during a 3-minute observation period. During this observation, each instance where a fish visibly consumed a food pellet was considered a feeding strike, aligning with the methodology outlined by O’Connor and Booth (52). In cases where a fish did not engage in feeding throughout the observation period, we recorded a bite rate of 0. Additionally, we noted the time elapsed until the fish-initiated feeding, which was set at 180 seconds for those fish that did not feed during the observation time.

(iii) **Shelter response** of the fish. A short (10cm) length of a cylindrical PVC pipe was introduced into the test tank, and the shelter response was categorized into different levels: -0: Fish exhibited no reaction to the structure and immediately swam away, maintaining normal swimming patterns. -1: Fish displayed no noticeable response to the structure and continued with their normal swimming patterns. -2: Fish exhibited no initial use of the structure for at least 60 seconds, but they eventually altered their normal swimming behaviours to approach and enter the structure. -3: Fish either immediately or very quickly (in less than 60 seconds) utilized the structure, modifying their normal swimming behaviours to approach and enter the shelter. These categories allowed us to rank and evaluate the shelter response of the fish under observation.

(iv) **Fish boldness,** a small block structure was introduced into the test tank with minimal disturbance to the fish. This block structure was chosen as it represents a novel object that doesn’t mimic any natural structure encountered by fish in their natural habitat, aligning with the methodology employed in prior experiments (56). Fish boldness was categorized into different levels: - 0: Fish immediately fled away from the structure, swimming in the opposite direction without any approach to the structure. -1: Fish exhibited no response to the structure, maintaining their normal swimming patterns, and did not approach the structure. - 2: Fish refrained from investigating the structure until at least 30 seconds into the observation period. They eventually altered their swimming behaviours to approach the structure, with a mean approach distance of 2-4 body lengths from the structure. -3: Fish immediately or very quickly (in less than 30 seconds) investigated the structure. They closely approached the structure and initiated physical contact, such as feeding strikes or bumping, at a distance of 1- 2 body lengths from the structure. These categories allowed us to assess and rank the boldness of the fish under observation based on their reactions to the introduced block structure.

(v) **Predator escape response** of the fish. A plastic fishing lure was employed as a proxy for a potential predator. This fishing lure was introduced into the test tank using a cable system, simulating a predator’s sudden appearance while minimizing the potential influence of human presence, in line with the methodologies outlined by Djurichkovic, Donelson (53) and Figueira, Curley (60).

(vi) **Burst-swimming speed** was calculated by measuring the distance (in centimetres) covered by the fish within the 2 seconds following the release of the fishing lure (61). The fish’s escape response was categorized into different levels: - 0: Fish displayed no response to the disturbance, maintaining normal behaviour both during and after the disturbance, with no alterations in swimming patterns. -1: Fish exhibited a slight delay in response to the disturbance (1-2 seconds). They initiated an escape response within 10 seconds but then quickly returned to their normal behaviour. During this response, they moderately increased their swimming speed and displayed erratic shifts in their swimming angle. -2: Like Category - 1, fish showed a slight delay in response (1-2 seconds). They initiated an escape response within 30 seconds and subsequently resumed normal behaviour. Their swimming speed moderately increased, and they exhibited erratic shifts in swimming angle. -3: Fish displayed an immediate or nearly immediate response to the disturbance. Their escape response continued for more than 30 seconds. This response was characterized by a significant increase in swimming speed, substantial erratic behaviour, and/or it was followed by freezing behaviour. These categories allowed for the evaluation and ranking of the fish’s escape responses in response to the simulated predator disturbance, offering insights into their predator avoidance behaviour.

(vii) **Fish metabolism** is difficult to measure and as such, oxygen consumption rate is used as a proxy (62). Resting metabolic rate (RMR) is evaluated by calculating the oxygen consumption of a fish while at rest (MO_2rest_), whereas the maximum metabolic rate (MMR) is estimated by calculating the maximum oxygen consumption (MO_2max_) when the fish is actively swimming. Aerobic scope was then calculated as the difference between the maximum and minimum oxygen consumption as per equation 3 from Measuring fish metabolism directly can be challenging, which is why oxygen consumption rate serves as a practical proxy, as described by McMahon, Parsons (62). In this context, two key metabolic parameters are assessed: - Resting Metabolic Rate (RMR): This metric is determined by measuring the oxygen consumption of a fish while it is at rest, denoted as MO_2rest_.

Maximum Metabolic Rate (MMR): The maximum metabolic rate is estimated by calculating the maximum oxygen consumption, referred to as MO_2max_, while the fish is actively swimming. Aerobic scope, which reflects the fish’s capacity for aerobic metabolism, is then calculated as the difference between the maximum and minimum oxygen consumption levels. This calculation is based on Equation 3 from McMahon, Parsons (62):

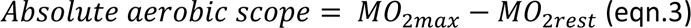

This enables evaluation of the range of oxygen consumption rates that a fish can sustain, providing insights into its metabolic capacity and energy utilization under different conditions. MO_2max_, MO_2rest_ and the absolute aerobic scope were quantified in mg O_2_ kg^-1^ h^-1^ as specified by the equation 4 from McMahon, Parsons (62):

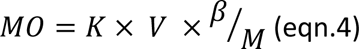

where *K* is the linear rate of decline (kPah^-1^) in the oxygen content over time (*h*) in the respirometry chamber, *V* is the volume of the chamber in L, *β* is the water solubility (depending on temperature and salinity, mg O_2_ L^-1^ kPa^-1^) and *M* is the fish’s mass (kg) (63).

The methodology for measuring oxygen use in *G. subfasciatus* and *P. sexlineatus* is based on the procedures described in Donelson et al. (2011) and McMahon et al. (2020), which involve intermittent respirometry techniques to monitor oxygen consumption rates in these fish species.

### **e)** Statistical analysis

To assess whether individual fish performance was preserved across temperatures, linear regressions were used to test the relationship between individual performance at 16°C and individual performance at 20°C (each data point consists of one individual’s performance at each of the two temperatures). A strongly significant positive result would indicate that performance at 16°C could be considered as a good predictor of performance at 20°C as expected, showing the relative performance among individuals is preserved to some degree across temperatures.

Linear regressions were used for instantaneous growth rate, change in total length, aerobic scope, bite rate and burst speed; while ordinal regression was used for shelter response, boldness, and escape response. First, the effect of food and order of temperatures treatment was assessed for each species and metric with ANOVAs. As food had no effect across all metrics, low food and high food treatment were combined for analysis. For the metrics that presented significance in the order of temperatures, separate regressions were run for the different order of temperatures groups (ascending/descending) (ANOVA results reported in supplementary Table 1).

Assumptions of normality and homogeneity of variances were assessed with the Shapiro-Wilk normality test and Levene’s Equality of Error Variance test, respectively. Any data that were found to violate the assumptions were log(x+1) or double-square root transformed, and outliers were removed following the interquartile range method (64). The data were statistically analysed with SPSS Statistics (65) using a significance level (α) of 0.05.

## 3. Results

Individual performance was generally not maintained across temperatures, that is, individual performance at 16°C did not generally predict individual performance at 20°C. A few exceptions were seen where performance was maintained across temperatures for *C.australis* boldness, *G.subfasciatus* aerobic scope and *P.sexlineatus* change in total length, bite rate, shelter and boldness. Table 2 summarises relationships between individual performance at 16°C vs 20°C for the performance metrics for the three species, with only 4 of 23 relationships being positive and 1 being negative (see summary Table 1).

**Table 1.**
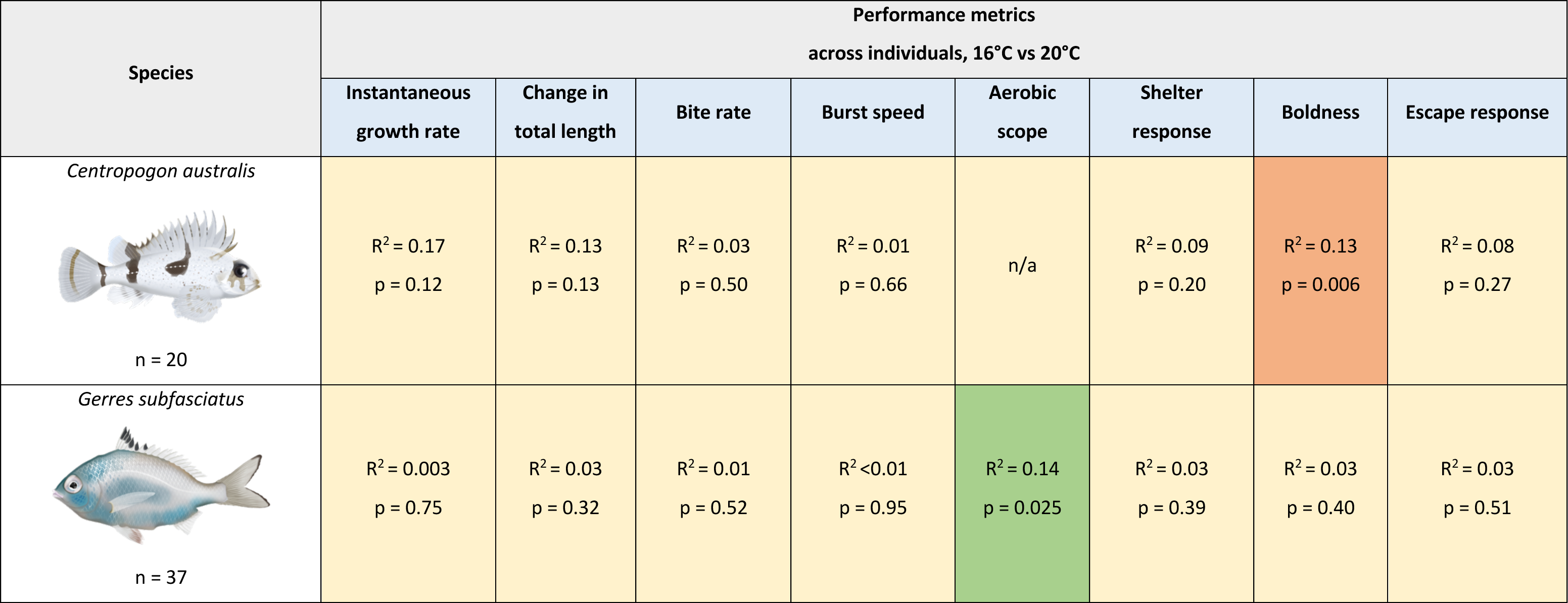
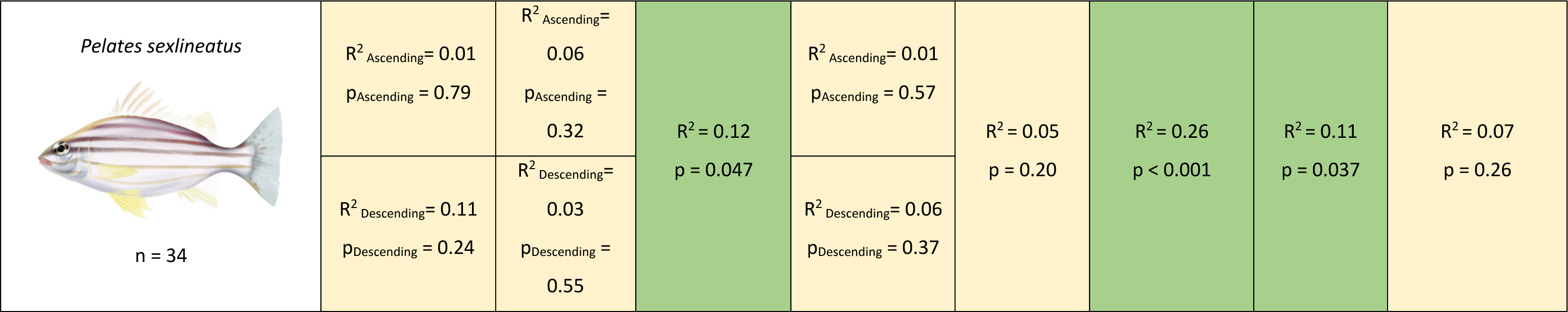
Relationship patterns in *Centropogon australis, Gerres subfasciatus* and *Pelates sexlineatus* individuals across physiological and behavioural performance (performance metric measured at 16°C vs performance metric measured at 20°C) with R^2^ and p-values. Separate regressions were conducted in case order of temperature was significant. ‘Ascending’ indicates ascending order of temperatures (16°C → 20°C) and ‘Descending’ indicates descending order of temperatures (20°C → 16°C). Red indicates a significantly negative relationship; green indicates a significantly positive relationship and yellow indicates the lack of a significant relationship. For linear regressions, Pearson’s R^2^ is reported, while for ordinal regressions, McFadden pseudo-R^2^ is reported.

| Species | Performance metrics<br>across individuals, 16°C vs 20°C |  |  |  |  |  |  |  |
| --- | --- | --- | --- | --- | --- | --- | --- | --- |
|  | Instantaneous growth rate | Change in total length | Bite rate | Burst speed | Aerobic scope | Shelter response | Boldness | Escape response |
| *Centropogon australis*<br>n = 20 | $R^2 = 0.17$<br>p = 0.12 | $R^2 = 0.13$<br>p = 0.13 | $R^2 = 0.03$<br>p = 0.50 | $R^2 = 0.01$<br>p = 0.66 | n/a | $R^2 = 0.09$<br>p = 0.20 | $R^2 = 0.13$<br>p = 0.006 | $R^2 = 0.08$<br>p = 0.27 |
| *Gerres subfasciatus*<br>n = 37 | $R^2 = 0.003$<br>p = 0.75 | $R^2 = 0.03$<br>p = 0.32 | $R^2 = 0.01$<br>p = 0.52 | $R^2 < 0.01$<br>p = 0.95 | $R^2 = 0.14$<br>p = 0.025 | $R^2 = 0.03$<br>p = 0.39 | $R^2 = 0.03$<br>p = 0.40 | $R^2 = 0.03$<br>p = 0.51 |
| *Pelates sexlineatus*<br>n = 34 | $R^2_{\text{Ascending}} = 0.01$<br>$p_{\text{Ascending}} = 0.79$ | $R^2_{\text{Ascending}} = 0.06$<br>$p_{\text{Ascending}} = 0.32$ | $R^2 = 0.12$<br>$p = 0.047$ | $R^2_{\text{Ascending}} = 0.01$<br>$p_{\text{Ascending}} = 0.57$ | $R^2 = 0.05$<br>$p = 0.20$ | $R^2 = 0.26$<br>$p < 0.001$ | $R^2 = 0.11$<br>$p = 0.037$ | $R^2 = 0.07$<br>$p = 0.26$ |
| | $R^2_{\text{Descending}} = 0.11$<br>$p_{\text{Descending}} = 0.24$ | $R^2_{\text{Descending}} = 0.03$<br>$p_{\text{Descending}} = 0.55$ | | $R^2_{\text{Descending}} = 0.06$<br>$p_{\text{Descending}} = 0.37$ | | | | |

In *C. australis,* a negative relationship was present between boldness at 16°C and at 20°C (χ^2^= 8.62, df= 3, *p* = 0.006; McFadden R^2^ = 0.13). In *G. subfasciatus*, a positive relationship was present between aerobic scope at 16°C and aerobic scope at 20°C (ANOVA, F_1,35_= 5.45, *p* = 0.025; R^2^ = 0.14), where aerobic scope at 16°C explained 14% of the variation in aerobic scope length at 20°C (F_1,35_ = 5.45, p-value= 0.025). In *P. sexlineatus*, a positive relationship was present between bite rate at 16°C and at 20°C (ANOVA, F_1,32_= 4.28, *p* = 0.047; R^2^ = 0.12), where performance was maintained across temperatures and bite rate at 16°C explained 12% of the variation in bite rate at 20°C (F_1,32_ = 4.28, p-value= 0.047). A positive relationship was present between shelter response at 16°C and at 20°C (χ^2^= 20.12, df= 2, *p* < 0.001; McFadden R^2^ = 0.26) and between boldness at 16°C and at 20°C (χ^2^= 8.50, df= 3, *p* = 0.037; McFadden R^2^ = 0.11).

Across all the other metrics, the relationship between individual performance at 16°C and performance at 20°C were non-significant. For instance, in *C. australis*, the instantaneous growth rate at 16°C did not reliably predict growth at 20°C (ANOVA, F_1,13_= 2.81, p = 0.12; R_2_ = 0.17). Similar trends were observed in the changes in total length at 16°C and 20°C (ANOVA, F_1,17=_ 2.55, p = 0.13; R_2_ = 0.13), bite rate (ANOVA, F_1,17_= 0.48, p = 0.50; R_2_ = 0.03), burst speed (ANOVA, F_1,14_= 0.20, p = 0.66; R_2_ = 0.01), shelter response at 16°C and 20°C (χ2= 4.62, df= 3, p = 0.20; McFadden R_2_ = 0.09), and escape response at 16°C and 20°C (χ2= 3.91, df= 3, p = 0.27; McFadden R_2_ = 0.08). In *G. subfasciatus*, the instantaneous growth rate at 16°C didn’t foresee growth at 20°C (ANOVA, F1,32= 0.10, p = 0.75; R2 = 0.003). Similar patterns were observed in the changes in total length at 16°C and 20°C (ANOVA, F_1,29_= 1.01, p = 0.32; R_2_ = 0.03), bite rate (ANOVA, F_1,35_= 0.42, p = 0.52; R_2_ = 0.01), burst speed (ANOVA, F_1,33_= 0.005, p = 0.95; R_2_ <0.01), shelter response (χ2= 3.01, df= 3, p = 0.39; McFadden R_2_ = 0.03), boldness (χ2= 2.94, df= 3, p = 0.40; McFadden R_2_ = 0.03), and escape response (χ2= 2.30, df= 3, p = 0.51; McFadden R_2_ = 0.03).

Likewise in *P. sexlineatus*, no significant relationship was present between instantaneous growth rate at 16°C and instantaneous growth rate at 20°C (ANOVA_Ascending Temperature_, F_1,15_= 0.08, *p* = 0.79; R^2^ = 0.01; ANOVA_Descending Temperature_, F_1,12_= 1.55, *p* = 0.24; R^2^ = 0.11). Analogously, a similar pattern was observed in change in total length across temperatures (ANOVA_Ascending Temperature_, F_1,16_= 1.03, *p* = 0.32; R^2^ = 0.06; ANOVA_Descending Temperature_, F_1,13_= 0.38, *p* = 0.55; R^2^ = 0.03), burst speed (ANOVA_Ascending Temperature_, F_1,14_= 0.34, *p* = 0.57; R^2^ = 0.01; ANOVA_Descending Temperature_, F_1,14_= 0.87, *p* = 0.37; R^2^ = 0.06),aerobic scope at (ANOVA, F_1,31_= 1.69, *p* = 0.20; R^2^ = 0.05) and escape response across temperatures (χ^2^= 4.02, df= 3, *p* = 0.26; McFadden R^2^ = 0.07).

Overall, there are considerable differences in the number of significant results among species, while *C. australis* and *G. subfasciatus* defied expectations with fewer significant relationships, *P. sexlineatus* largely followed anticipated patterns, showing both negative and positive relationships.

## 4. Discussion

Contrary to expectations, this study highlighted differences in individual response to temperature, where surprisingly individual relative performance was not maintained across temperatures for most performance metrics. Some fish performed relatively better at lower temperatures, others had moderate performance across both temperatures, and others had higher performance the high temperature. This suggests that, for each species tested, performance was maintained across temperatures for some variables, while for most variables “niche performance” differed .

The individuals that had similar activity and growth rates across both temperatures therefore presented high contextual plasticity, thus a wider thermal optimum and broader tolerance to temperature change. In contrast, the individuals that had markedly greater growth and activity rates when exposed to high temperatures showed low contextual plasticity (20). These patterns are obscured in most population studies reporting overall means and variances (13). If we had ignored individual variability and only assessed each fish at a particular temperature, the behavioural estimates would have been biased by the temperature itself (20). For instance, since fish tend to exhibit bolder behaviours in warmer temperatures, an individual that is usually shy might seem bold, even bolder than a naturally bold fish in colder conditions when observed in warm environments. Considering plasticity, an individual with high contextual plasticity could have appeared relatively bold at cooler temperatures. In contrast, the same individual at a warmer temperature would be identified as shy compared to an individual with low contextual plasticity, where the changes in boldness with temperature are much sharper (20).

In contrast with our finding that performance was mostly not preserved across treatments, studies that assessed the Amazon molly (*Poecilia formosa*) and the green swordtail (*Xiphophorus helleri*) dominance showed that individuals maintained their losing/winning performance over time (15). Individuals that only experienced winning interactions in the early life stages held the same trend later in life, remaining top-ranked individuals in the hierarchy while losers remained at the bottom rank and neutral individuals were in the middle (15, 17). Additionally, a study on the spotted catshark (*Scyliorhinus canicula)* demonstrated how shark social network position was consistent and maintained across different habitats (16). This study highlighted how previous social experiences could have substantial and long-lasting consequences on the adult population’s social behaviour and structure (15).

Across all species, growth rate and change in total length at 16°C were not significant predictors of relative performance at 20°C. The magnitude of individual variability is surprising where not all individuals followed the population pattern with an increase in performance with temperature, but some moved from low to high performance in cold temperatures while others maintained average performance at both temperatures. This highlights the strong individual differences in thermal optima (6). As aforementioned, when below the thermal optimum, an increase in temperature usually provides larger aerobic scope and, thus, more energy available for growth which would explain the higher growth rates at higher temperatures experienced by most individuals (11). Colder waters lower the energetic costs due to activity that could allow the fish to invest in growth, if a sufficient aerobic scope is available (66). Thus, this could justify the higher growth rates in some fish at 16°C.

Differences in growth rates with temperature have important implications for fish shoals, as fish that grow faster become larger, affecting social hierarchy (67). Larger fish are usually more aggressive and dominant members that outcompete the suburbanites in access to food and mates (67, 68). This social hierarchy can impact the social dynamics within a group, such as reproductive success and therefore affect future generation genetics (69). Additionally, this increases the homogenisation of reaction responses while lowering population resilience as the genetic pool shrinks (70). This study highlights that these social hierarchies could be modified by a temperature increase as the individuals that grow the most (and thus are likely to be dominant) at 16°C would not be the same ones that grow the most at 20°C. A study on the African cichlid fish species, *A. burtoni* found that size differences that were once thought to be negligible (<10% body length) provided a substantial advantage to dominant and larger individuals (71). While fish were housed individually in our study for logistics reasons, future work should explore our predictions against direct social interactions across temperature treatments.

However, despite the fact that faster-growth-rate and larger individuals benefit from this attribute in the social hierarchy, the size-selection that operates at a fisheries level can act against a rapid growth rate, where larger individuals are more likely to be selected and removed (18). Additionally, warm adapted-faster-growing genotypes might be at a disadvantage in a winter situation, because the more robust appetite and, thus, higher feeding rates that are required to sustain the higher growth rates might induce more risk-taking and bolder behaviours to forage and thus increase the risk of encounter fishing gear and vulnerability to it (18). A lake experiment showed that genotypes that grow faster were captured at three times the rate of the slow-growing ones (18).

*G.subfasciatus* aerobic scope at 16°C was a good predictor of aerobic scope at 20°C, which was expected as these temperatures are within the fish thermal range, and a slight increase in temperature below thermal optimum is expected to boost metabolism (72, 73). This might result from the consistency in sub-cellular and cellular components and processes within individuals (74). Additionally, it has been shown that resting metabolic rate is consistently different (repeatable) across individuals in birds, mammals and some fish (75–80) and thus we would expect such consistency as environmental conditions are modified. Additionally, relationships between metabolic rates at different temperatures could be influenced by the difference in individual responses to stress that fish with a shy or bold phenotype experience during the actual metabolic test. For instance, shy individuals have been shown to increase their oxygen consumption during respirometry due to their higher sensitivity to confinement and handling stress (81).

However, individual *P.sexlineatus’* aerobic scope at the lower temperature was not a good predictor of its aerobic scope with a temperature increase (82). The processes behind intraspecific variation in metabolic rate are uncertain (13). Intraspecific variation in metabolic rate could be the physiology of individuals, such as the leakiness of mitochondrial membranes or the protein turnover rates (83). Some studies showed how variation in body-size correlated with metabolic rate, revealing a positive relationship between metabolic rate and the size of metabolically costly organs such as the brain and heart (84). In *Salmo trutta*, intraspecific changes in metabolic rate were positively associated with the activity of some mitochondrial enzymes (85), while changes in erythrocyte size were found to be negatively correlated with metabolic rate in the loach *Cobitis taenia* (86). These species differences in aerobic scope associations across temperatures highlight how performance maintenance, such as aerobic scope, is mainly species and individual-specific. Therefore different individuals and species will be affected differently by climate change (52).

Activity such as boldness was previously identified as an intrinsic trait across individuals (87). Members of the same species frequently behave differently, some being more aggressive or bold and others being more docile or shy (83), with some individuals having an innate propensity to be more active all the time (87). As such, this might justify the relationship between boldness at 16°C and 20°C in *P.sexlineatus* and *C.australis* as fish adapted to the changes in temperature to maintain the same relative performance.

*P. sexlineatus* exhibited a significant relationship across temperature treatments for bite rate, boldness, and shelter response. Under natural conditions, an individual’s food intake rates and, thus, growth can be heightened through a combination of boldness and activity (20). And this process could therefore increase the positive relationships in these metrics across temperatures where a consistent increase in behavioural activity such as boldness can allow for higher foraging efficiency thus, higher food intakes (88).

Skeletal muscle activity affects individual performance in essential activities such as locomotion, prey capture and predator avoidance (89). The lack of relationship in escape response across temperatures detected in all species could be motivated by differences in cellular physiology that act at the base of responsiveness to stimuli such as predators (90).

The difference in swimming performance across individuals influences their spatial positioning within a school. Individuals with a relatively high aerobic scope and gait transition speed might locate themselves in anterior positions within the school (Metcalfe et al. 2016). These higher swimming speeds would allow these individuals to be found at the front of the school while performing other tasks such as feeding or digestion (13). Consequently, these intrinsic differences in swimming capabilities may be associated with differences in intra-species migration success and probability of evading predator attacks (91, 92). Individual differences in boldness positively influence migratory inclination, the possibility of becoming dominant, foraging success, and reproductive performance. Bold actions, however, also have adverse fitness effects, such as greater predation risk, which can lower long-term survival (23).

Differences in shelter usage across individuals might be linked to trade-offs between energy usage and foraging success (34). For instance, sheltering might lower fish energy expenditure linked to performing mechanical tasks (e.g., swimming) (93). Sheltering can also lower the energy expenditure of non-mechanical tasks such as thermoregulation (94), or high-energy activities such as camouflage, alertness and vigilance (95). Therefore individual differences in shelter use might have significant impacts on individual survival and growth (34).

In addition to physiological traits such as metabolism, different activity types can induce fish to choose different strategies to cope with stressors such as temperature changes. This is performed by selecting and migrating to cold habitats that lower energy expenditure (less active fish) or warmer habitats that promote a more active lifestyle (66).

The processes mentioned above highlight behavioural traits’ influence on the fish stock’s social hierarchy, and where metrics are related to social dominance for example, it may be a mechanism by which the dynamics of estuarine fish populations could differ under climate change. Thus, the lack of relationship between many of these performance metrics at the lower vs higher temperature indicates that stock structures will likely be modified if temperatures increases, possibly lowering fisheries resilience (70). Individuals that currently present a physiological advantage at lower temperatures might drop to lower subordinate levels in favour of the individuals that are better adapted to these new warmer environments (18). As individuals present different performance optima, these individuals performance niches may serve as a buffer to population-level responses, initially resulting in a reshuffling of individual performance ranks as temperature increases, without altering the population average. After this reshuffling is finalized, a detectable population response may emerge as the temperature continues to rise. Thus, individual performance niches have the potential to mitigate population responses, or at the very least, impede our ability to perceive them.

## 5. Conclusion

Individual fish exhibited substantial differences in performance across temperatures. This highlights how climate change will likely favour a niche of adapted individuals who will thrive in warmer conditions. In contrast, the subset of individuals that perform best at lower temperatures might encounter higher competition as their niche temperature habitat retreats, which could pose a risk for them to transfer their ’cold acclimated’ hereditary traits (13). Thus, future communities will likely present different genetic compositions, physiological and behavioural characteristics than the current ones. The underlying shift in genetic composition, to genotypes that perform better under warmer temperatures, may actually mask a climate change response from studies that continue to focus on population averages when testing performance; changes in relative individual performance might initially counterbalance a population-level response, buffering climate change responses and potentially hindering our ability to detect them. Detecting individual changes in performance may provide an early warning for future population-level consequences of warming. To further enhance the knowledge of individual fish performance related to climate change, future studies could include broader performance metrics (such as courtship, shoaling aggression, and impact of temperature on neurological systems) (4, 96) and more fish species (97).

## Ethics

All practices in this study were conducted following the NSW Department of Primary Industries scientific collection permit (Permit No: F94/696(A)-9.0) and the University of Technology Sydney Animal Ethics protocols (Permit No: ETH21- 6609).

## Data Accessibility

The data are provided in electronic supplementary material.

## Declaration of AI use

The authors declared that we have not used AI-assisted technologies in creating this article.

## Author’s contributions

All authors contributed to data generation, analysis, manuscript preparation, and conceptualizing the paper’s structure.

## Supporting information

Bellotto et al Supplementary Data

## Acknowledgments

Our thanks to UTS for support and facilities. Dr. J. Donelson and Dr. W. Figueira provided vital guidance and equipment. Special thanks to lab staff: Helen Price, Rachel Keeys, Scott Allchin, and Sue Fenech for their assistance.

