## Supplementary material for "Individual performance niches may buffer population responses to climate change in estuarine fishes": Bellotto et al Supplementary Data

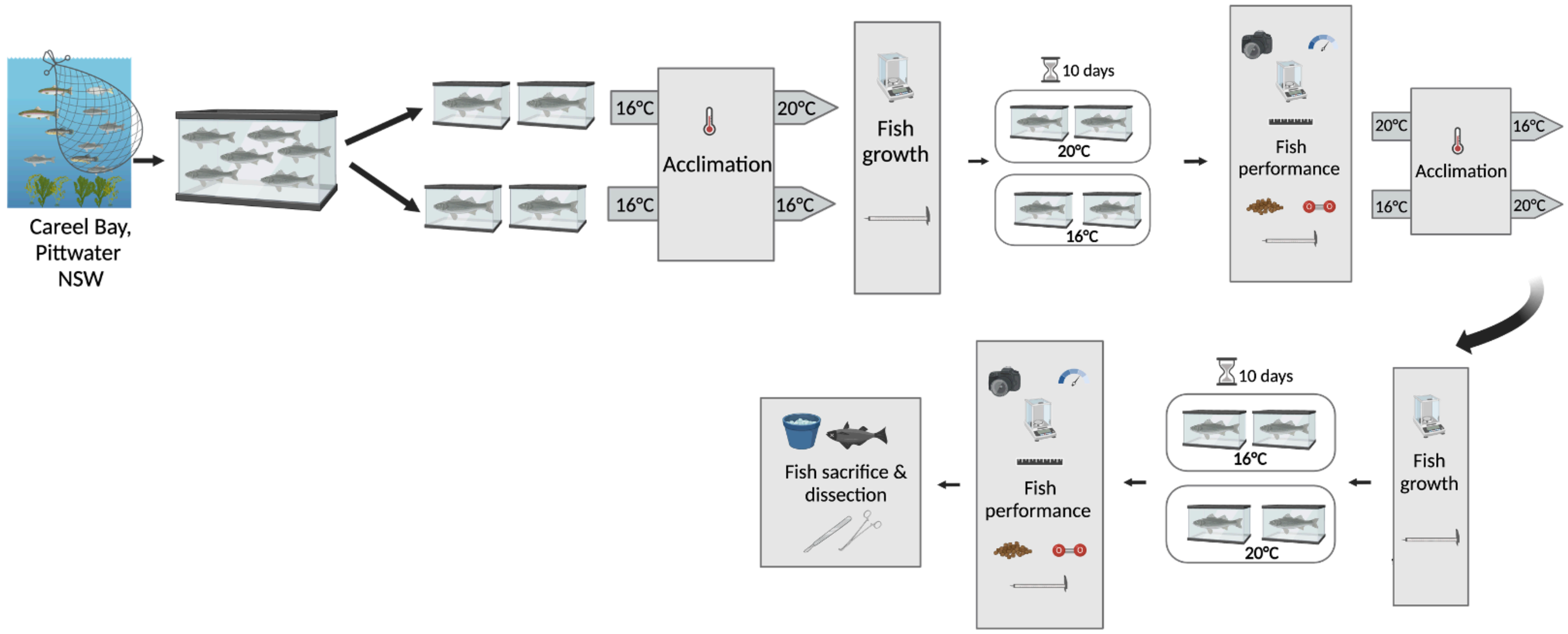
Careel Bay,
Pittwater
NSW
16°C
16°C
Acclimation
20°C
16°C
Fish
growth
10 days
20°C
16°C
Fish
performance
20°C
16°C
Acclimation
16°C
20°C
Fish
growth
10 days
16°C
20°C
Fish
performance
Fish sacrifice &
dissection

**(A)**

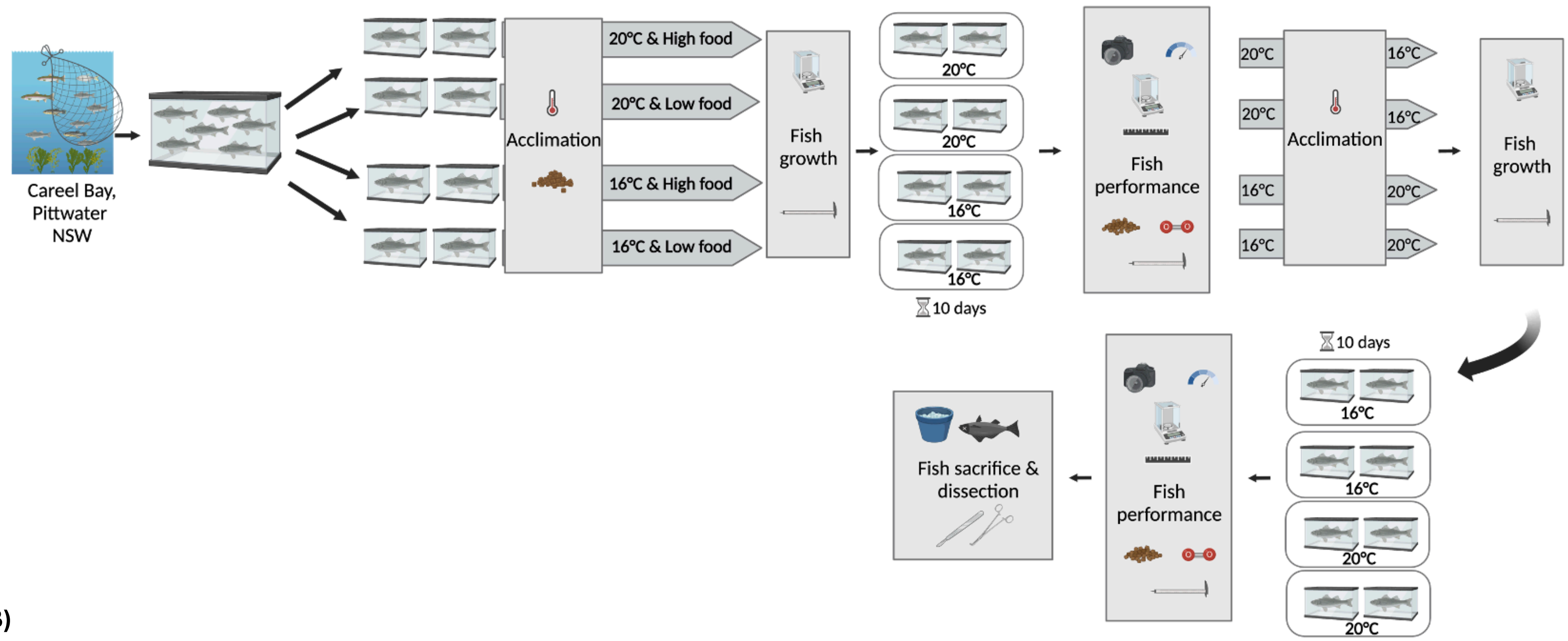


**(B)**

**Figure S1. (A)** Diagram of experimental design for the eastern fortescues, *Centropogon australis* (n = 20), with temperature (16°C vs 20°C) as factor. **(B)** Diagram of experimental design for the common silver biddy *Gerres subfasciatus* (n=37) and the eastern striped trumpeter, *Pelates sexlineatus* (n=34) with temperature (16°C vs 20°C) and food (low vs high food regime) as factors. A separate experiment was conducted for each species. Low food regimes were fed 0.5% fish weight in fish pellets, High food regimes were fed 1% fish weight in fish pellets. Created with BioRender (2023).

**Table S1.** ANOVA (F values, degrees of freedom, p-values) results for the effect of food regime and order of temperatures treatment across species.

| Species | Performance metric | Factor | F | df | p-value |
|---|---|---|---|---|---|
| ***Centropogon australis*** | Instantaneous growth rate | Order of temperatures | 2.061 | 1,33 | 0.161 |
| | Change in total length | Order of temperatures | 3.414 | 1,35 | 0.074 |
| | Bite rate | Order of temperatures | 0.137 | 1,35 | 0.714 |
| | Burst speed | Order of temperatures | 0.048 | 1,32 | 0.828 |
| | Shelter response | Order of temperatures | 0.421 | 1,36 | 0.521 |
| | Boldness | Order of temperatures | 1.574 | 1,29 | 0.218 |
| | Escape response | Order of temperatures | 0.928 | 1,36 | 0.342 |
| ***Gerres* subfasciatus** | Instantaneous growth rate | Food regime | 0.260 | 1,63 | 0.612 |
| | | Order of temperatures | 2.674 | 1,63 | 0.107 |
| | Change in total length | Food regime | 0.132 | 1,58 | 0.717 |
| | | Order of temperatures | 0.705 | 1,58 | 0.404 |
| | Bite rate | Food regime | 0.631 | 1,66 | 0.430 |
| | | Order of temperatures | 2.128 | 1,66 | 0.149 |
| | Burst speed | Food regime | 0.026 | 1,64 | 0.873 |
| | | Order of temperatures | 0.288 | 1,64 | 0.594 |
| | Aerobic scope | Food regime | 0.009 | 1,64 | 0.926 |
| | | Order of temperatures | 0.667 | 1,64 | 0.417 |
| | Shelter response | Food regime | 1.730 | 1,66 | 0.163 |
| | | Order of temperatures | 1.922 | 1,66 | 0.170 |
| | Boldness | Food regime | 1.380 | 1,66 | 0.244 |

| | | | | | |
|---|---|---|---|---|---|
| | | Order of temperatures | 0.451 | 1,66 | 0.504 |
| | Escape response | Food regime | 0.004 | 1,66 | 0.949 |
| | | Order of temperatures | 1.817 | 1,66 | 0.182 |
| ***Pelates sexlineatus*** | Instantaneous growth rate | Food regime | 0.713 | 1,57 | 0.402 |
| | | Order of temperatures | 12.533 | 1,57 | <0.001 |
| | Change in total length | Food regime | 0.276 | 1,59 | 0.601 |
| | | Order of temperatures | 14.860 | 1,59 | <0.001 |
| | Bite rate | Food regime | 1.073 | 1,60 | 0.304 |
| | | Order of temperatures | 0.074 | 1,60 | 0.786 |
| | Burst speed | Food regime | 0.568 | 1,58 | 0.454 |
| | | Order of temperatures | 16.454 | 1,58 | <0.001 |
| | Aerobic scope | Food regime | 0.035 | 1,59 | 0.851 |
| | | Order of temperatures | 0.409 | 1,59 | 0.525 |
| | Shelter response | Food regime | 0.016 | 1,60 | 0.901 |
| | | Order of temperatures | 0.200 | 1,60 | 0.656 |
| | Boldness | Food regime | 1.540 | 1,60 | 0.220 |
| | | Order of temperatures | 0.341 | 1,60 | 0.561 |
| | Escape response | Food regime | 0.501 | 1,60 | 0.482 |
| | | Order of temperatures | 0.557 | 1,60 | 0.458 |
